# Uncovering parasite diversity in Ecuadorian wildlife: new trypanosomatid species and novel reservoir hosts for *Leishmania amazonensis*

**DOI:** 10.64898/2026.06.22.733918

**Authors:** Manuel Alejandro Coba-Males, Alexei Yu. Kostygov, Henry D. Naranjo, Jaime A. Salas, Sandra Enríquez, Pablo Medrano-Vizcaíno, David Brito-Zapata, Sofía Ocaña-Mayorga, Juan Carlos Navarro, Jazzmin Arrivillaga, Sarah Martin-Solano, Gabriel Alberto Carrillo-Bilbao, Wilmer Narváez, Manuela González-Suárez, Vyacheslav Yurchenko, Ana Poveda

## Abstract

Wildlife hosts play important roles in the ecology and transmission of vector-borne parasites, yet information on host associations remains scarce in many biodiverse tropical regions. Within a One Health framework, characterizing parasite diversity in wildlife can improve understanding of ecosystem health and disease emergence. Road-killed animals provide a non-invasive opportunity to investigate host–parasite interactions while minimizing disturbance to natural habitats.

We screened 127 liver and intestinal tissue samples obtained from 76 road-killed vertebrates collected near protected areas in two Ecuadorian biodiversity hotspots, the Tropical Andes and Chocó-Darién, for trypanosomatids and other vector-borne microorganisms. Molecular analyses targeted the 18S rRNA and cytochrome b genes of trypanosomatids and included additional screening for *Trypanosoma cruzi*, *Trypanosoma rangeli*, *Rickettsia* spp., and piroplasmids. Twenty-nine samples were positive for kinetoplastids. We detected diverse trypanosomatids representing the genera *Leishmania*, *Porcisia*, *Trypanosoma*, *Phytomonas*, *Blastocrithidia*, and *Obscuromonas*, as well as free-living kinetoplastids of the order Neobodonida. The most frequently detected species was *Leishmania amazonensis*, identified in 17 samples from at least 13 species of birds, reptiles, and caecilians, predominantly in liver tissue, suggesting previously unrecognized host associations. We also identified a putatively novel species of *Porcisia* and three potentially undescribed avian trypanosomes belonging to the subgenus *Ornithotrypanum*. No evidence of *T. cruzi*, *T. rangeli*, *Rickettsia* spp., or piroplasmids was found.

Our findings identify birds, reptiles, and caecilians as potential reservoir hosts of *L. amazonensis.* In addition, we substantially expanded current knowledge of kinetoplastid diversity in Ecuadorian wildlife. This study demonstrates the value of road-killed animals as a practical, non-invasive resource for wildlife pathogen surveillance and highlights the importance of integrating biodiversity research into One Health approaches to better understand parasite transmission dynamics in rapidly changing tropical ecosystems.

**Author summary:** Many parasites that affect humans circulate naturally in wildlife, but identifying their animal hosts is often difficult in remote, biodiverse regions. We used road-killed animals as a non-invasive source of biological material to investigate parasites in wildlife from two biodiversity hotspots in Ecuador. By analyzing tissues from birds, reptiles, amphibians, and mammals, we found a remarkable diversity of kinetoplastid flagellates, a group that includes the agents of Chagas disease and leishmaniasis. Although we did not detect human-infective trypanosomes, we repeatedly identified *Leishmania amazonensis* (a species causing human disease) in birds, reptiles, and caecilians. These vertebrate groups have not previously been recognized as potential hosts of this parasite. We also discovered several undescribed trypanosomatid species, emphasizing how little is known about parasite diversity in tropical wildlife. Our results show that road-killed animals can provide valuable information on host–parasite interactions without disturbing living populations. Such surveillance contributes to One Health efforts by improving our understanding of how environmental change, wildlife, and human health are interconnected.

## Introduction

Within the One Health framework, coordinated improvement of human, animal, and ecosystem health critically depends on comprehensive characterization of biodiversity, including host-associated communities [1–3]. In this context, identifying parasites infecting wildlife, which often serve as a reservoir of human pathogens, is important both for assessing health of ecosystems and for addressing the medical aspect of One Health research. However, collecting such data can be particularly challenging in remote regions with limited funding and research infrastructure. A practical solution is the analysis of the road-killed animals, a non-invasive approach that avoids entering natural habitats or handling live specimens, thereby minimizing impact on ecosystems [4,5]. Roads and human-modified landscapes constitute ecological interfaces, where humans, domestic animals, wildlife, and vectors increasingly interact, enabling pathogen spillover and emergence [6].

The family Trypanosomatidae is a diverse and speciose group of parasitic flagellates, encompassing two categories of species: monoxenous, which develop in a single (typically insect) host, and dixenous, which alternate between a vertebrate or plant host and an insect vector [7–9]. Although most studies have understandably focused on *Leishmania* and *Trypanosoma* that are pathogenic to humans, pets, and livestock, interest is growing in flagellates that parasitize other animals, abundant in natural habitats and important for ecosystem functioning [10–12].

*Trypanosoma* is the largest genus within the family Trypanosomatidae, currently encompassing over 500 described species and parasitizing all classes of vertebrates [13]. In Ecuador, as in the rest of Latin America, the most important trypanosome species is *T. cruzi*, the etiological agent of the Chagas disease transmitted by triatomine bugs [14,15]. Numerous vertebrate wildlife species have been implicated as reservoirs for this parasite [16]. Another trypanosome, *T. rangeli* has the same vectors and geographic distribution as *T. cruzi*, and can be confused with it in microscopical and serological tests [17–19]. While generally regarded as non-pathogenic to humans, *T. rangeli* is important from the epidemiological perspective [19]. It has been documented in Ecuador, although the number of reports is limited [20,21].

*Leishmania* is a genus uniting multiple species and attracting most research attention in many regions of the world [22]. These parasites cause leishmaniasis, a disease with three main clinical manifestations: cutaneous, mucocutaneous, and visceral, of which only the former two have been reported in Ecuador with prevalence reaching thousands of cases annually [23,24]. At least seven species of *Leishmania* (subgenera *Leishmania* and *Viannia*), namely *L. amazonensis*, *L. braziliensis*, *L. guyanensis*, *L. lainsoni*, *L. mexicana*, *L. naiffi*, and *L. panamensis*, vectored by four sand fly species, have been identified and implicated in human infections in this country [25,26]. However, very little is known about animal hosts, either domestic or wild, for *Leishmania* in Ecuador [27]. This knowledge gap is particularly relevant in developing countries, where accelerated environmental change, road expansion, and biodiversity disturbance may alter host–vector–pathogen interactions. Meanwhile, there is growing evidence that wild animals, not limited to mammals, can serve as reservoir hosts for *Leishmania* pathogenic to humans [22,28–31].

So far, in Ecuador, trypanosomatids have been intensively sampled in insects [32–36], while records from the wild vertebrates are scarce [37]. In this work, we investigated road-killed vertebrates collected in the vicinity of several protected areas in Ecuador for the presence of trypanosomatids and some other vector-borne microorganisms of medical and veterinary relevance. Our molecular screening documented multiple monoxenous and dixenous trypanosomatid species, most of which were not previously recorded. In addition, we identified potential novel reservoir hosts, such as birds, reptiles, and caecilians (amphibians), for *Leishmania amazonensis*. These findings expand the knowledge of parasitic flagellates, including species that infect humans, and highlight the value of road-killed animals sampling for more detailed investigations into the diversity and ecology of these fascinating parasites.

## Materials and methods

### Road-kill sampling

The sampling was conducted in areas within two biodiversity hotspots in Ecuador (Fig 1). In the tropical Andes (Amazonian province Napo), 240 km of roads passing by four protected areas (Antisana Ecological Reserve, Sumaco-Napo-Galeras National Park, Cayambe-Coca National Park, Colonso Chalupas Biological Reserve, and LLanganates National Park) were surveyed from November 2020 to March 2021. In Chocó-Darien moist forests (coastal provinces of Guayas and Santa Elena), specimens were collected between July 2021 and February 2023 along 200 km of roads near four conservation areas (Manglares El Salado Wildlife Production Reserve, Cerro Blanco Protected Forest, Parque Lago National Recreation Area, and Manglares El Morro wildlife refuge). Details of the collection procedures have been described previously [38]. The work was performed under permissions MAAE-ARSFC-2020-0791 and MAAE-ARSFC-2021-1862 (for collection of biological material) and MAAE-DBI-CM-2021-0215 (for molecular analyses and access to genetic resources) from the Ministry of Environment, Water, and Ecological Transition of Ecuador.

**Fig 1.**
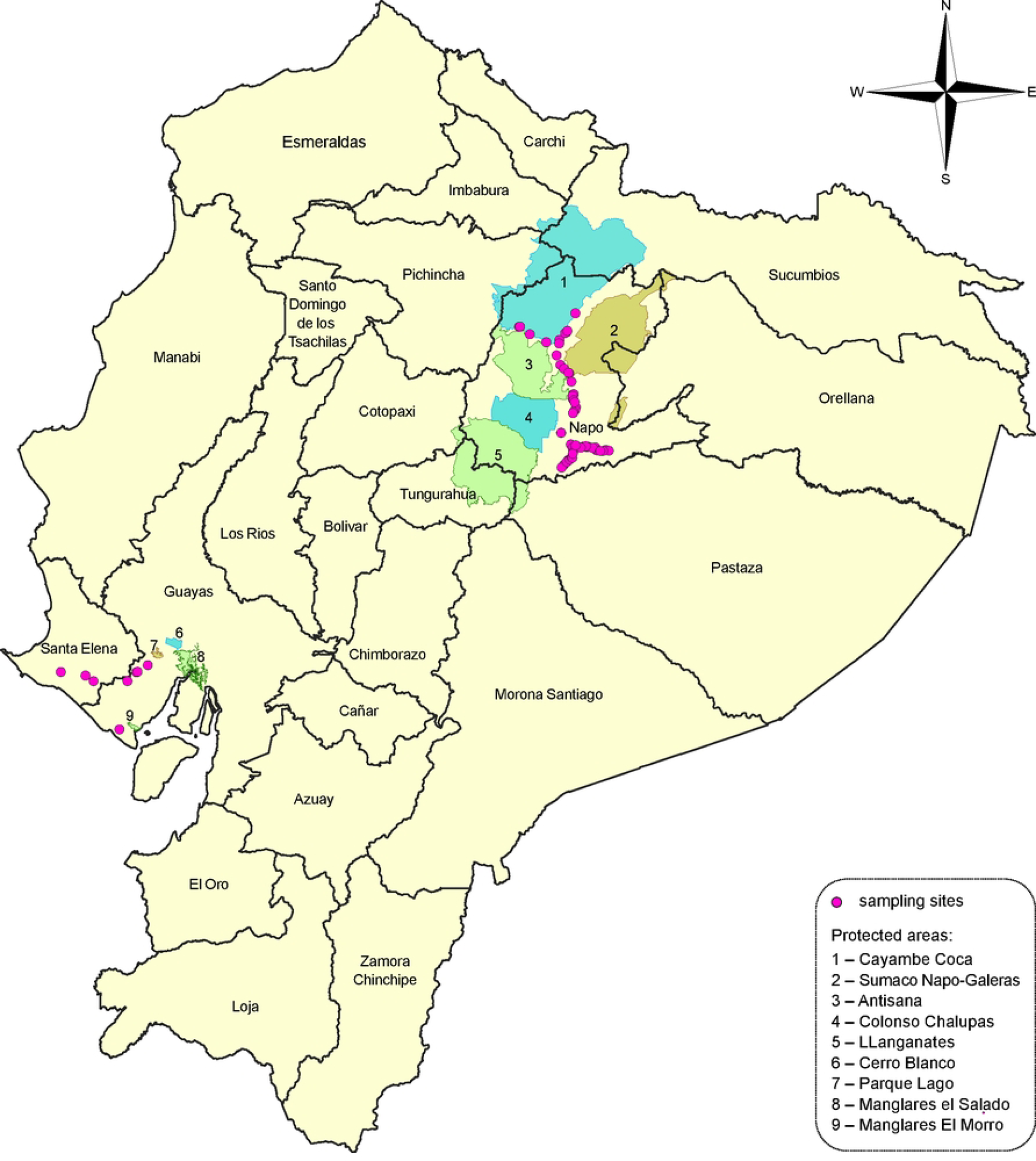
Map of Ecuador showing sampling sites. Provinces are labeled directly on the map. Protected areas (in cyan, green, and olive) are marked by numbers explained in the graphical legend.

When the condition of a carcass permitted, liver and intestine were sampled. A total of 127 tissue samples (109 from the tropical Andes and 18 from Chocó-Darien) were collected across 77 individuals, including 3 caecilians, 27 birds, 6 mammals, and 41 reptiles (Table 1, S1 Table).

**Table 1.** Summary of collected samples from road-kill animals.

| Organ | Tropical Andes |  |  |  |
| --- | --- | --- | --- | --- |
|  | Amphibians | Birds | Mammals | Reptiles |
| Intestine | 2 | 21 | 2 | 28 |
| Liver | 2 | 19 | 3 | 32 |
| Subtotal | 4 | 40 | 5 | 60 |
| Chocó-Darién |  |  |  |  |
| Intestine | 0 | 4 | 1 | 2 |
| Liver | 0 | 5 | 2 | 4 |
| Subtotal | 0 | 9 | 3 | 6 |
| **Total** | **4** | **49** | **8** | **66** |

### Molecular analyses

DNA from the organ samples (∼ 25 mg each) was isolated using the PureLink Genomic DNA Mini Kit (Thermo Fisher Scientific, Waltham, USA) following the manufacturer’s instructions. The presence of trypanosomatids was initially assessed by nested PCR of the 18S rRNA gene. DNA samples from cultured *L. amazonensis*, *L. mexicana*, and *L. tarentolae*, whose identity had been confirmed by sequencing the 18S rRNA and *cytb* genes, served as positive controls. The first round used primers SLF (5′-GCTTGTTTCAAGGACTTAGC-3′) and S762R (5′-GACTTTTGCTTCCTCTAATG-3′), followed by the second round with primers S825F (5′-CGAACAACTGCCCTATCAGC-3′) and SLIR (5′-GACTACAATGGTCTCTAATC-3′), yielding a product of approximately 950 bp [39,40]. For positive samples, an overlapping ∼900 bp fragment was amplified using primers S823 (5′-ACCGTTTCGGCTTTTGTTGG-3′) and S662R (5′-ACATTGTAGTGCGCGTGTC-3′) [41]. Assembly of the two fragments produced a approximately 1.5 kb sequence, representing about 70% of the total gene length.

The cytochrome *b* (*cytb*) gene was amplified by nested PCR as in [24,42], using the direct primer 397F (5′-AGCGGAGAGGAAAGAAAAG-3′) along with two reverse primers 404R1 (5′-CTACAATAAACAAATCATAATATACAAT-3′) and 405R2 (5′-CTACAATAAACAAATCATAATATGCAAT-3′) in the first round. For the second round, the direct primer 399F (5′-GTGTAGGTTTTAGTTTAGG-3′) along with two reverse primers 406R (5′-GTTCACAATAAAATGCAAAT-3′) and 407R2 (5′-GTTCGCAATAAAATGCAAAT-3′) were used, yielding a ∼ 600 bp product.

For all samples, the presence of *T. cruzi* and *T. rangeli* was additionally assessed by amplification of the conserved domain of kinetoplast minicircles using primers 121 (5′-AAATAATGTACGGGKGAGATGCATGA-3′) and 122 (5′-GGTTCGATTGGGGTTGGTGTAATATA-3′), which for these species are expected to yield bands in the range of 300-400 bp [43].

To test for the presence of *Rickettsia* spp., the *gltA* gene was targeted with previously described primers CS-78 (5′-GCAAGTATCGGTGAGGATGTAAT-3′), CS-323 (5′-GCTTCCTTAAAATTCAATAAATCAGGAT-3′), and EHR16SR (5’-TAGCACTCATCGTTTACAGC-3’) [44]. Screening for piroplasmids (*Babesia* spp.) was performed *via* amplification of the 18S rRNA gene using primers BAB 143-167 (5′-CCGTGCTAATTGTAGGGCTAATACA-3′) and BAB 694-667 (5′-GCTTGAAACACTCTARTTTTCTCAAAG-3′), as described previously [45].

All positive amplicons were sequenced at Macrogen (Seoul, South Korea), and the sequences obtained were submitted to GenBank; accession numbers are listed in S1 Table.

### Phylogenetic analyses

The sequences of 18S rRNA and cytochrome b (*cytb*) genes obtained in this study were combined with those available in the core nucleotide and whole genome shotgun sequence databases of GenBank or genomic database of TriTrypDB [46]. For *Leishmania*, additional sequences were assembled from NCBI sequence read archives (SRAs) using reference-based approach in Geneious Prime v. 2026.0.2 as described previously [47]. Five datasets were created: i) concatenated 18S rRNA + *cytb* genes for dixenous Leishmaniatae (i.e., genera *Leishmania*, *Endotrypanum*, and *Porcisia*); ii) 18S rRNA gene for bird-associated *Trypanosoma* (i.e. subgenera *Avitrypanum*, *Ornithotrypanum* and *Trypanomorpha*) combined with closely related (ingroup) *Megatrypanum*, as well as *Schizotrypanum* as an outgroup; iii) 18S rRNA gene for Blastocrithidiinae; iv) 18S rRNA gene for *Phytomonas*, with *Herpetomonas* and *Lafontella* included as an outgroup; and v) 18S rRNA gene for Neobodonida, with Parabodonida as an outgroup. The sequences were aligned using MAFFT v. 7.490 [48] using E-INS-i and L-INS-I algorithms for 18S rRNA and *cytb* gene sequences, respectively.

Maximum likelihood (ML) analysis was conducted in IQ-TREE v. 3.0.1 [49] under automatically selected models with edge support assessed by standard bootstrap method with 1,000 replicates. For dixenous Leishmaniatae, an edge-proportional partition model (accounting for the differences between the two genes) was applied to the concatenated dataset. An additional analysis was performed for the subset including only *cytb* sequences, as this gene allows better phylogenetic resolution, which may be masked by conflicts with the 18S rRNA gene, which is less informative for this group.

Bayesian inference was performed in MrBayes v. 3.2.7 [50] under the models mirroring those in ML analysis, but with substitution matrices generalized to GTR and unlinking branch lengths in the partitioned model for the concatenated dataset. The analysis was run for 1,000,000 generations, sampling frequency of 100 and other parameters set to default.

## Results

Of the total 127 samples, only 29 tested positive by nested PCR for the 18S rRNA gene of Kinetoplastea (Fig 2, S1 Table). Among the obtained sequences, 26 corresponded to trypanosomatids of the genera *Leishmania*, *Porcisia*, *Trypanosoma*, *Phytomonas*, *Blastocrithidia*, and *Obscuromonas*, while the remaining three belonged to kinetoplastean flagellates of the order Neobodonida, which were also amplified by the primers used (Fig 2, S1 Table).

**Fig 2.**
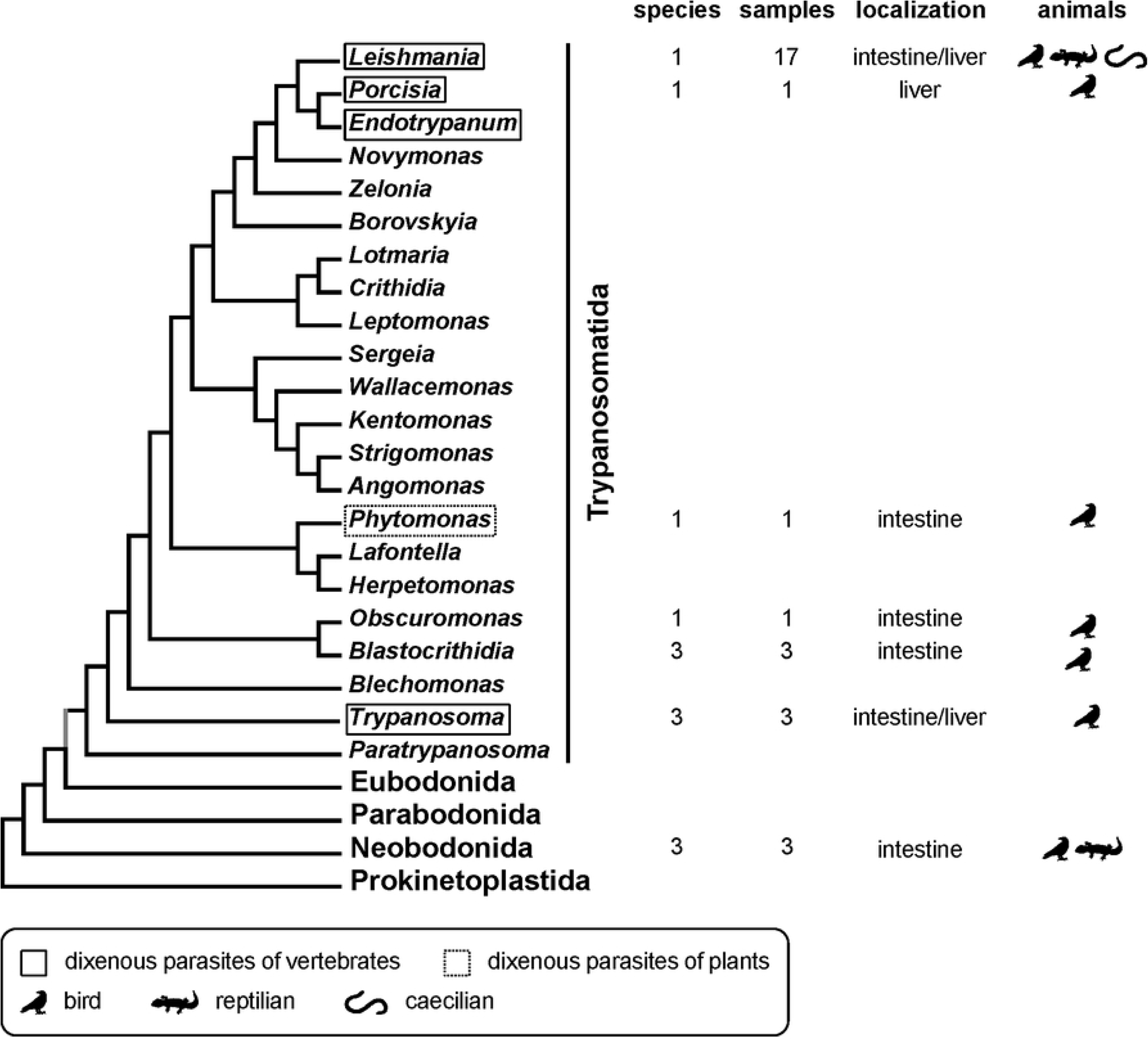
Summary of samples positive for Kinetoplastea, showing their taxonomic distribution, species counts, localization, and vertebrate group. The cladogram is based on previously published phylogenetic inferences for Kinetoplastea [51] and Trypanosomatidae [7].

The screening for other parasites (*Rickettsia* and *Babesia* spp.) did not reveal any positives. Similar negative results were obtained when testing for the presence of *T. cruzi* and *T. rangeli* with minicircle-targeting primers (Table S1). This is in line with the analysis based on the 18S rRNA gene, which also did not detect these two species of trypanosomes.

The most prevalent detected trypanosomatids belonged to a single species of *Leishmania*, *L. amazonensis*, documented (either by 18S rRNA or by combination of 18S rRNA and *cytb* genes) in 17 samples across at least 13 species of birds, caecilians, and reptiles (Figs 2 and 3). All but one of these 17 samples originated from locations in the Amazonian province Napo, Tropical Andes (S1 Table). All obtained sequences of the two genes were identical to the reference sequences from GenBank, suggesting that all identified parasites belonged to a single strain. The majority (12/17) of the samples positive for *L. amazonensis* (and at least one from each vertebrate group) derived from liver, suggesting that non-specific detection of *Leishmania* in the analyzed animals is not likely. A whooping motmot *Momotus subrufescens* (Momotidae, Coraciiformes; sample 971_ECCD) was found to harbor a member of the related genus *Porcisia* in the liver, again suggesting a specific infection. The phylogenetic analysis revealed that this trypanosomatid represents a previously undescribed species of this poorly studied and enigmatic taxon (Fig 3).

**Fig 3.**
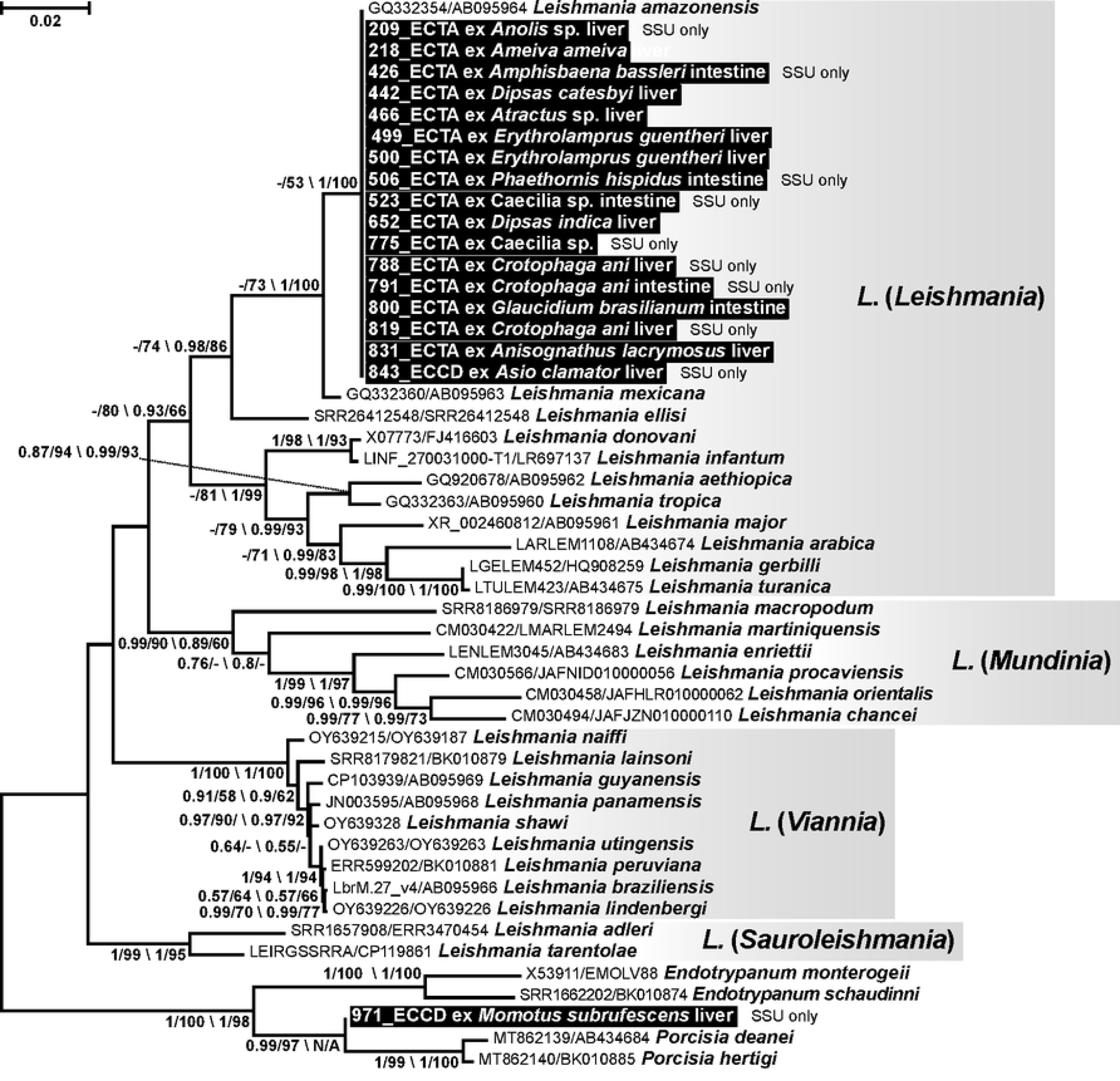
Maximum likelihood tree of dixenous Leishmaniatae based on concatenated *cytochrome b* and 18S rRNA gene sequences. The tree is rooted according to a previous phylogenomic inference [52]. The new isolates are highlighted in black. Numbers at branches represent Bayesian posterior probabilities and bootstrap supports, with respective values below 0.5 and 50% replaced with dashes. These values are shown as two backslash-separated sets: for the concatenated dataset and for *cytochrome b* only. The scale bar denotes the number of substitutions per site.

While our analyses did not identify any human-parasitic trypanosomes, we documented presence of avian *Trypanosoma* spp. of the subgenus *Ornithotrypanum* in samples from birds (Fig 4). A parasite found in the intestine of a black-billed thrush *Turdus ignobilis* (Turdidae, Passeriformes) belonged to a clade recently recovered in a study of trypanosome fauna in Colombia [53]. All other sequences in this clade originated from blood of various birds of the order Passeriformes: i) crimson-backed tanager *Ramphocelus dimidiatus* (Thraupidae), ii) common tody-flycatcher *Todirostrum cinereum* (Tyrannidae), iii) saffron finch *Sicalis flaveola* (Thraupidae), iv) great kiskadee *Pitangus sulphuratus* (Tyrannidae), and v) olivaceous greenlet *Hylophilus olivaceus* (Vireonidae) [53]. Considering strong association of trypanosomes in this clade with passerine birds, the positive specimen from *T. ignobilis* (728_ECTA) could represent a specific infection. However, the omnivorous habit of this bird species [54], along with the detection of the trypanosome only in the intestinal (and not in the liver) sample does not allow ruling out the possibility that the sequence originated from the intestinal content, which could include an infected insect vector.

**Fig 4.**
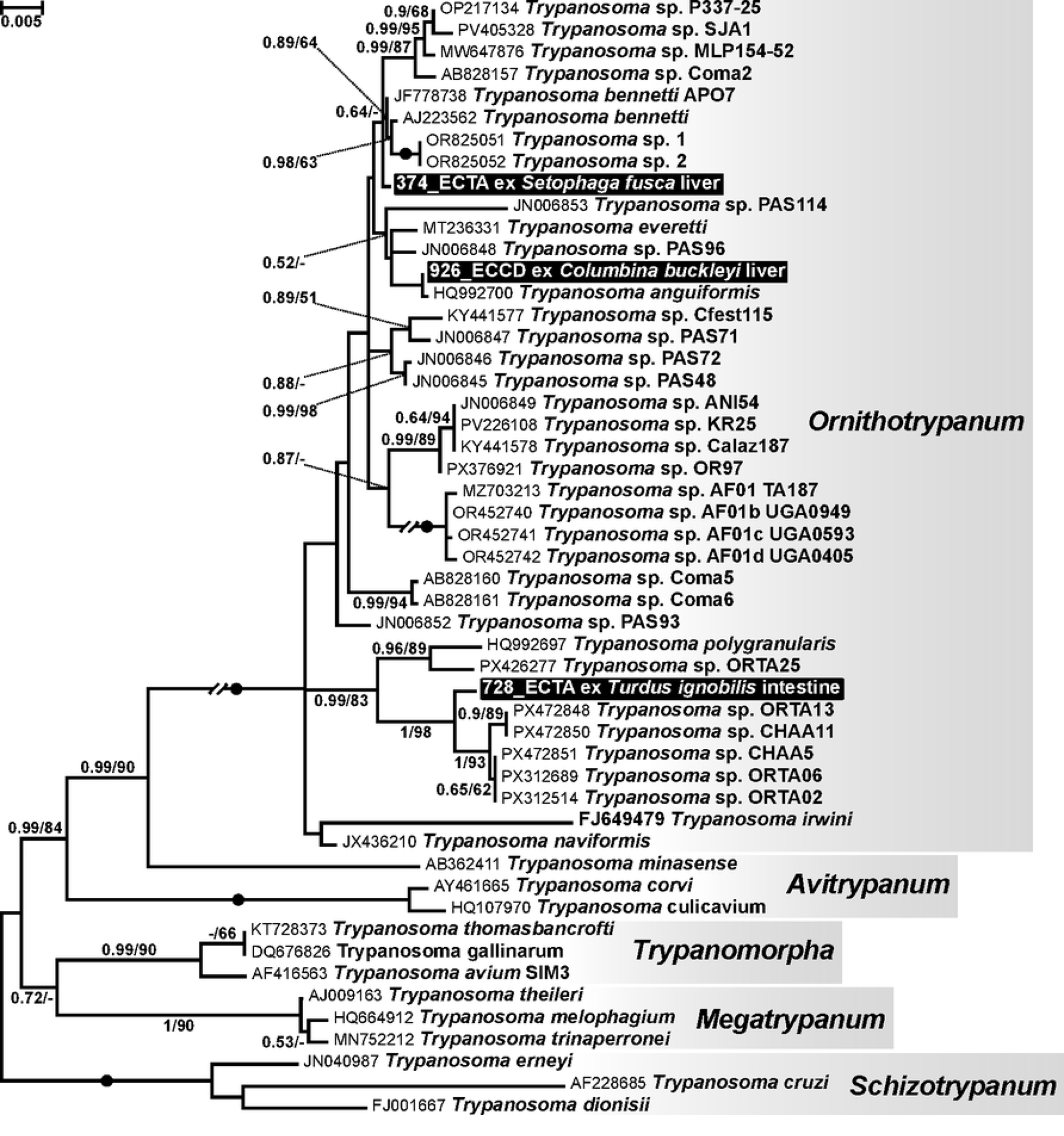
Maximum likelihood tree of bird-associated *Trypanosoma* spp. based on 18S rRNA gene sequences. The tree focuses on the subgenus *Ornithotrypanum*, in which new isolates were detected (highlighted in black), while other subgenera are included for the phylogenetic context. Sequences of *Schizotrypanum* were used for rooting in accordance with previously published phylogenomic analysis [7]. Double-crossed branches are shown at 50% of their original length. Numbers at branches indicate Bayesian posterior probabilities and bootstrap supports; values below 0.5 and 50%, respectively, are replaced with dashes, and maximal support (1/100) is indicated by filled circles. The scale bar denotes the number of substitutions per site.

Two other detected trypanosome sequences fell into a part of the tree, where high content of short sequences significantly reduced phylogenetic resolution, not allowing to clearly determine affiliations of the flagellates. A trypanosome from the liver of a Ecuadorian ground dove *Columbina buckleyi* (Columbidae, Columbiformes; sample 926_ECCD) was very closely related to *T. anguiformes* described from the African olive sunbird *Cyanomitra olivacea* (Nectariniidae, Passeriformes) in Cameroon [55]. The differences in sequences were minimal, however, considering their incompleteness (1.5 and 0.8 kb) and incomplete overlap, we cannot assert whether they really belong to the same species. To our knowledge, *Ornithotrypanum* has previously been reported from a limited number of avian hosts, including Neotropical passerines, but not from Columbiformes [56]. A sequence of a trypanosome from the liver of a blackburnian warbler *Setophaga fusca* (Parulidae, Passeriformes, sample 374_ECTA) could not be properly positioned on the tree, as in addition to low phylogenetic resolution, no closely related sequences were available.

All other kinetoplastean species detected in this work originated from intestinal specimens. In four samples, representatives of the subfamily Blastocrithidiinae were documented, specifically, three *Blastocrithidia* spp. and one *Obscuromonas* sp. (Fig 5). One *Blastocrithidia* sp. from the common pauraque *Nyctidromus albicollis* (Caprimulgidae, Caprimulgiformes, sample 919_ECCD) had 18S rRNA sequence identical to that detected in the Cuban true bug *Paromius longulus* (Rhyparochromidae, Heteroptera) [57]. The two other *Blastocrithidia* spp. from the smooth-billed ani *Crotophaga ani* (Cuculidae, Cuculiformes, samples 644_ECTA, 660_ECTA) did not match any previous records. The only *Obscuromonas* sp. sequence was revealed in the gut of abovementioned *Setophaga fusca* (sample 375_ECTA) and was closely related to the one from *Dolichomiris linearis* (Miridae, Heteroptera) collected in Cuba [57]. Blastocrithidiinae are restricted to true bugs and, moreover, many of these flagellates are host species-specific [8,58]. Therefore, detection of such parasites in the gut of exclusively or predominantly insectivorous birds [59–61] is unlikely to indicate specific infection. Blastocrithidiinae form cyst-like amastigotes, which are extremely resistant to adverse environmental conditions [62]. This facilitates their survival in the gut content increasing chances of their detection.

**Fig 5.**
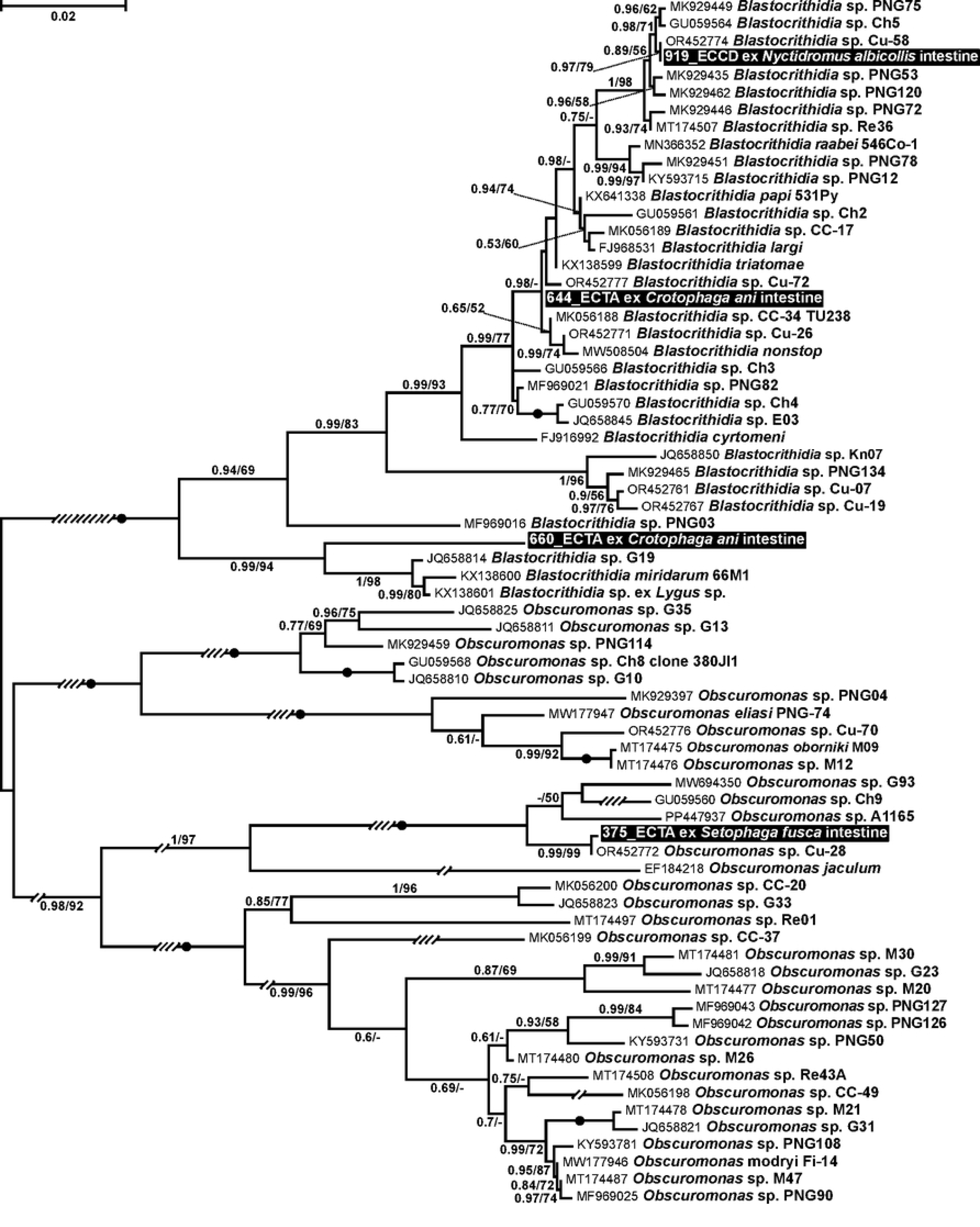
Maximum likelihood tree of Blastocrithidiinae based on 18S rRNA gene sequences. The tree is shown without an explicit outgroup and is rooted according to previously inferred relationships within the subfamily [8,58]. Sequences detected in this study are highlighted in black. Oblique lines crossing branches indicate the fold reduction of their original length. Numbers at branches represent Bayesian posterior probabilities and bootstrap supports; values below 0.5 and 50%, respectively, are replaced with dashes, and maximal support (1/100) is indicated by filled circles. The scale bar denotes the number of substitutions per site.

A single gut specimen of *Columbina buckleyi* (sample 929_ECCD) (the same individual, where a trypanosome was revealed) contained DNA of an unnamed *Phytomonas* sp., which was not closely related to any described species of this flagellate genus (Fig 6). The long-tailed mockingbird feeds on insects and fruits [63], both of which can be parasitized by phytomonads [64]. The dispersal stage of the life cycle of *Phytomonas* spp., the endomastigotes, are tailored to sustain a wide range of environmental conditions [62], again, ensuring better preservation of their DNA even in the decaying material.

**Fig 6.**
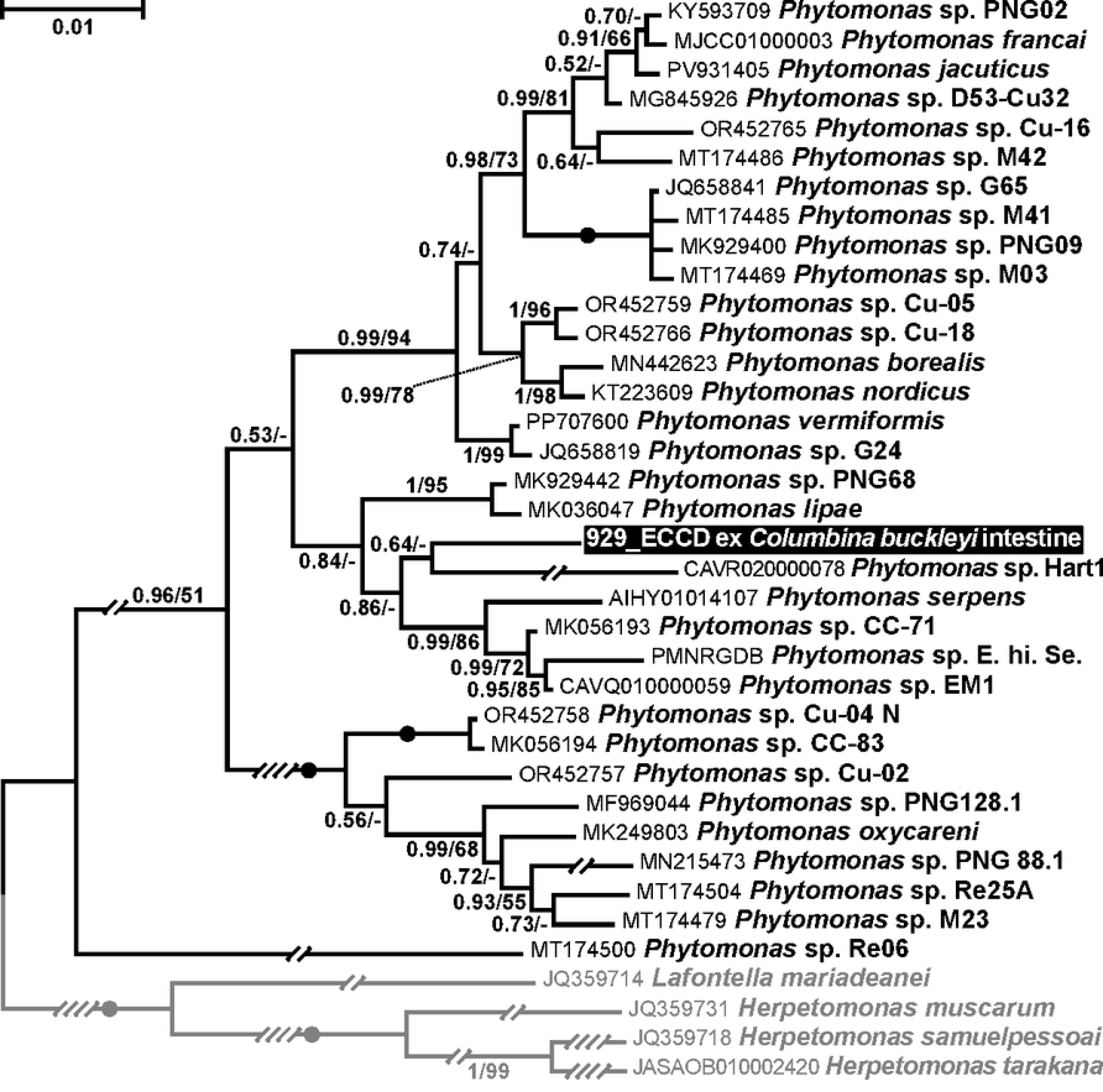
Bayesian tree of *Phytomonas* spp. based on 18S rRNA gene sequences. The tree is rooted with the sequences of *Herpetomonas* and *Lafontella* (shown in grey). Sequences detected in this study are highlighted in black. Oblique lines crossing branches indicate the fold reduction of their original length. Numbers at branches represent Bayesian posterior probabilities and bootstrap supports; values below 0.5 and 50%, respectively, are replaced with dashes, and maximal support (1/100) is indicated by filled circles. The scale bar denotes the number of substitutions per site.

Three sequences obtained in this work clustered among free-living kinetoplasteans of the order Neobodonida. The flagellates in two samples from colubrid snakes (52_ECTA and 318_ECTA, S1 Table) affiliated with the family Rhynchomonadidae (Fig 7). One of them was closely related to *Dimastigella* sp. (labeled as “*Phanerobia pelophila*” in the original publication [65]), while another represented a previously unrecognized lineage sister to all *Dimastigella* reported thus far. The third flagellate species detected in the intestine of *Momotus subrufescens* (sample 965_ECCD, the same individual as above harboring *Porcisia* sp.) fell into the paraphyletic family Neobodonidae, in which the majority of species were classified as the type species *Neobodo designis*. The detection of free-living kinetoplasteans in the intestine is not surprising since many of them form resistant cysts [8]. This has been documented for *Dimastigella* spp. [66], while the situation with *N. designis*-like flagellate is obscure.

**Fig 7.**
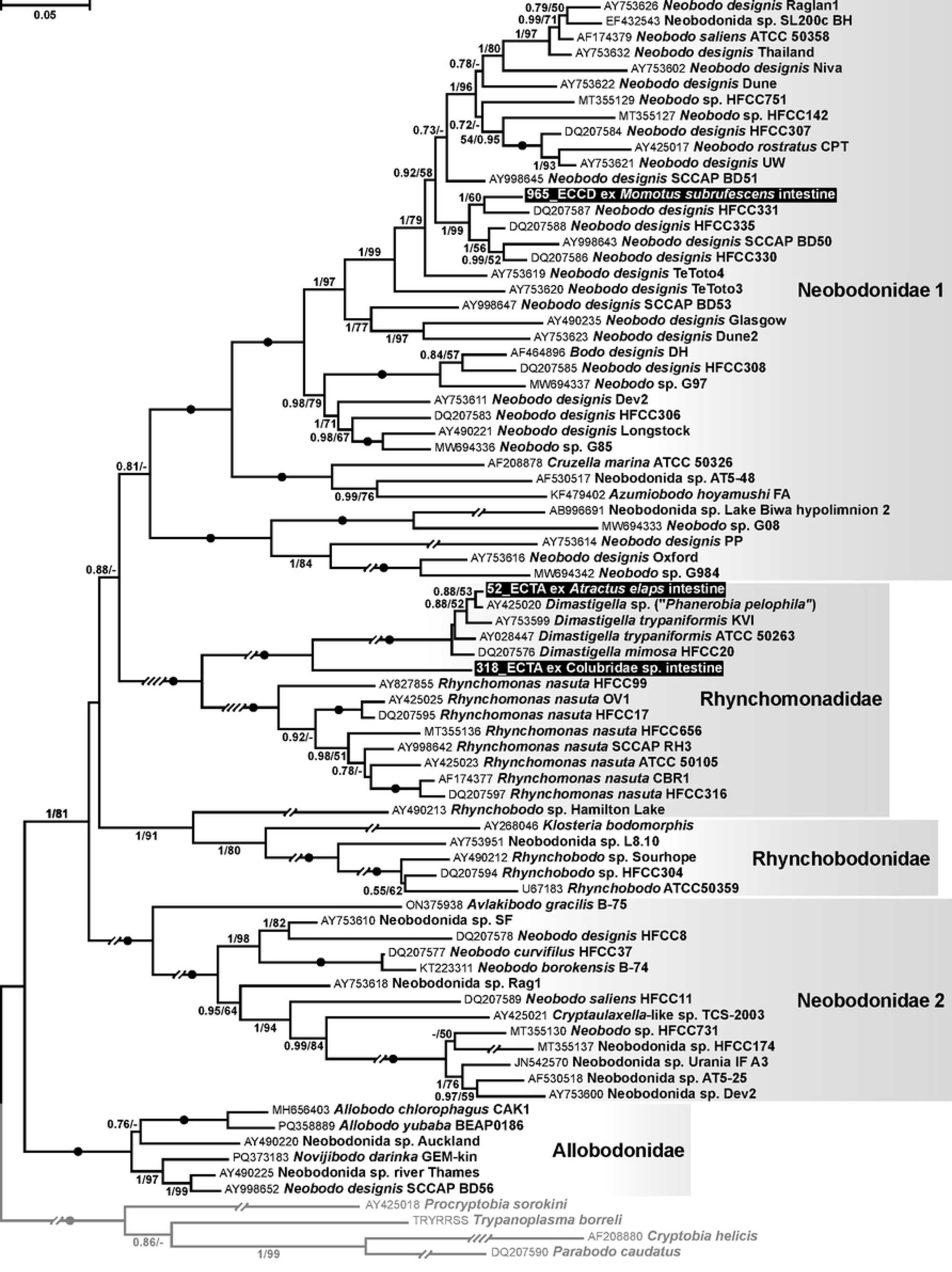
Maximum likelihood tree of Neobodonida based on 18S rRNA gene sequences. The tree is rooted with the sequences of Parabodonida (shown in grey). Sequences detected in this study are highlighted in black. Oblique lines crossing branches indicate the fold reduction of their original length. Numbers at branches represent Bayesian posterior probabilities and bootstrap supports; values below 0.5 and 50%, respectively, are replaced with dashes, and maximal support (1/100) is indicated by filled circles. The scale bar denotes the number of substitutions per site.

## Discussion

Here, we surveyed wildlife in two biodiversity hotspots of Ecuador for several disease-associated microorganisms in road-killed mammals, birds, reptiles, and amphibians using molecular methods. The screening revealed only diverse kinetoplastid flagellates. While no *T. cruzi* or *T. rangeli* infections were documented, many samples contained another human parasite, *L. amazonensis*. This allowed to expand the range of potential reservoir hosts for this important pathogen. To the best of our knowledge, only mammals have been implicated in this role for *L. amazonensis* [67]. Nonetheless, other human-infecting *Leishmania* spp., such as *L. infantum* and *L. tropica*, were recorded in the blood of birds and lizards [30,68–72]. One recently published study reported the presence of an unidentified species of *Leishmania* (for which human infectivity cannot be ascertained) in the blood of the toad *Rhinella horribilis* [73]. It must be noted that PCR detection of parasites, even in blood or such internal organs as liver, does not guarantee genuine infection. Trypanosomatid DNA can persist in blood for prolonged time after non-specific acquisition [74].

The presented study expands known diversity of Trypanosomatidae. Our screening revealed three potentially novel species of avian trypanosomes (subgenus *Ornithotrypanum*) in passerine and columbid birds, identifying previously unknown host-parasite systems. Here, the likelihood of infections’ specificity is higher than in the cases of *L. amazonensis* because the association of related trypanosomes with Passeriformes has been documented repeatedly in the past [53,55,75,76]. Considering that this group of trypanosomes remains poorly studied, any new data on their distribution are important.

Analysis of intestinal specimens in this work followed the same logic as in the studies using fecal samples: this allows detection of potential host-parasite associations though accompanied by a high background of non-specific material consumed with food [77]. Nonetheless, it cannot be excluded that in some instances, positivity of intestinal samples reflected a genuine infection. This concerns *Leishmania* and *Trypanosoma* spp., as in both cases the same or related parasites were detected in liver. We are convinced that the other identified kinetoplasteans (monoxenous *Blastocrithidia* and *Obscuromonas*, dixenous plant-infective *Phytomonas* spp., as well as non-parasitic neobodonids) likely represent passive content of the intestine. However, analysis of such samples gives insights into diversity of trypanosomatid fauna in a given ecosystem. In line with this, a wide range of monoxenous trypanosomatids and phytomonads has been documented in fecal samples of great apes [78].

Our pioneer study of Ecuadorian wildlife for the presence of human-parasitic trypanosomatids expanded screening to previously neglected groups of vertebrates and revealed potential novel reservoir hosts of *L. amazonensis*. These findings reinforce the importance of wildlife surveillance in tropical ecosystems for understanding the diversity of parasites and their hosts, which is a critical component of One Health strategies linking environmental integrity, animal health, and public health preparedness [79]. Besides facilitating identification of transmission networks within ecosystems for medically relevant parasites, our study advances the knowledge of the diversity of those trypanosomatids that have been long considered unimportant (monoxenous genera and *Phytomonas*) but have gained research attention in the last few decades.

The detection of *L. amazonensis* across multiple vertebrate groups suggests complex ecological transmission patterns that warrant further investigation from a One Health and eco-epidemiological perspective. Anthropogenic pressures, such as habitat fragmentation, road expansion, and increase of wildlife–human interfaces, influence parasite circulation dynamics. This highlights the need for interdisciplinary approaches integrating ecology, epidemiology, conservation biology, and public health.

## Data availability statement

All sequences obtained have been submitted to GenBank, with accession numbers listed in S1 Table.

## Supplementary data legends

**S1 Table.** Detailed information on collected specimens, including taxonomy of animals, sampled organs, collection localities, estimated time since death, and positivity for screened parasites. Due to the condition of the carcasses, some species could not be identified below family level. Some data were not collected during sampling.

## References

1. WHO (2025) The WHO One Health Initiative (OHI).

2. Marselle MR, Hartig T, Cox DTC, de Bell S, Knapp S, Lindley S, et al. Pathways linking biodiversity to human health: a conceptual framework. Environ Int. 2021;150.

3. Fagre AC, Cohen LE, Eskew EA, Farrell M, Glennon E, Joseph MB, et al. Assessing the risk of human-to-wildlife pathogen transmission for conservation and public health. Ecol Lett. 2022;25: 1534–1549.

4. Szekeres S, Docters van Leeuwen A, Toth E, Majoros G, Sprong H, Foldvari G Road-killed mammals provide insight into tick-borne bacterial pathogen communities within urban habitats. Transbound Emerg Dis. 2019;66: 277–286.

5. Coba-Males MA, Diaz M, Molina CA, Medrano-Vizcaino P, Brito-Zapata D, Martin-Solano S, et al. Gut bacterial communities in roadkill animals: a pioneering study of two species in the Amazon region in Ecuador. PLoS One. 2024;19: e0313263.

6. Caron A, Morand S, de Garine-Wichatitsky M (2012) Epidemiological interaction at the wildlife/livestock/human interface: can we anticipate emerging infectious diseases in their hotspots? A framework for understanding emerging diseases processes in their hot spots. In: Morand S, Beaudeau F, Cabaret J, editors. New frontiers of molecular epidemiology of infectious diseases. Dordrecht: Springer. pp. 311–332.

7. Kostygov AY, Albanaz ATS, Butenko A, Gerasimov ES, Lukeš J, Yurchenko V Phylogenetic framework to explore trait evolution in Trypanosomatidae. Trends Parasitol. 2024;40: 96–99.

8. Kostygov AY, Karnkowska A, Votýpka J, Tashyreva D, Maciszewski K, Yurchenko V, et al. Euglenozoa: taxonomy, diversity and ecology, symbioses and viruses. Open Biol. 2021;11: 200407.

9. Maslov DA, Opperdoes FR, Kostygov AY, Hashimi H, Lukeš J, Yurchenko V Recent advances in trypanosomatid research: genome organization, expression, metabolism, taxonomy and evolution. Parasitology. 2019;146: 1–27.

10. Maslov DA, Votýpka J, Yurchenko V, Lukeš J Diversity and phylogeny of insect trypanosomatids: all that is hidden shall be revealed. Trends Parasitol. 2013;29: 43–52.

11. Lukeš J, Butenko A, Hashimi H, Maslov DA, Votýpka J, Yurchenko V Trypanosomatids are much more than just trypanosomes: clues from the expanded family tree. Trends Parasitol. 2018;34: 466–480.

12. Lukeš J, Skalický T, Týč J, Votýpka J, Yurchenko V Evolution of parasitism in kinetoplastid flagellates. Mol Biochem Parasitol. 2014;195: 115–122.

13. Spodareva VV, Grybchuk-Ieremenko A, Losev A, Votýpka J, Lukeš J, Yurchenko V, et al. Diversity and evolution of anuran trypanosomes: insights from the study of European species. Parasit Vectors. 2018;11: 447.

14. Cucunubá ZM, Gutiérrez-Romero SA, Ramírez JD, Velásquez-Ortiz N, Ceccarelli S, Parra-Henao G, et al. The epidemiology of Chagas disease in the Americas. Lancet Reg Health Am. 2024;37: 100881.

15. Calvopina M, Segovia G, Cevallos W, Vicuña Y, Costales JA, Guevara A Fatal acute Chagas disease by *Trypanosoma cruzi* DTU TcI, Ecuador. BMC Infect Dis. 2020;20: 143.

16. Ibarra-Cerdeña CN, Valiente-Banuet L, Sanchez-Cordero V, Stephens CR, Ramsey JM *Trypanosoma cruzi* reservoir-triatomine vector co-occurrence networks reveal meta-community effects by synanthropic mammals on geographic dispersal. PeerJ. 2017;5: e3152.

17. Guhl F, Hudson L, Marinkelle CJ, Jaramillo CA, Bridge D Clinical *Trypanosoma rangeli* infection as a complication of Chagas’ disease. Parasitology. 1987;94: 475–484.

18. de Moraes MH, Guarneri AA, Girardi FP, Rodrigues JB, Eger I, Tyler KM, et al. Different serological cross-reactivity of *Trypanosoma rangeli* forms in Trypanosoma cruzi-infected patients sera. Parasit Vectors. 2008;1: 20.

19. Bayão TS, Cupertino MDC, Mayers NAJ, Siqueira-Batista R A systematic review of the diagnostic aspects and use of *Trypanosoma rangeli* as an immunogen for *Trypanosoma cruzi* infection. Rev Soc Bras Med Trop. 2020;53: e20190608.

20. Ocaña-Mayorga S, Aguirre-Villacis F, Pinto CM, Vallejo GA, Grijalva MJ Prevalence, genetic characterization, and 18S small subunit ribosomal RNA diversity of *Trypanosoma rangeli* in triatomine and mammal hosts in endemic areas for Chagas disease in Ecuador. Vector Borne Zoonotic Dis. 2015;15: 732–742.

21. Maiguashca Sánchez J, Sueto SOB, Schwabl P, Grijalva MJ, Llewellyn MS, Costales JA Remarkable genetic diversity of *Trypanosoma cruzi* and *Trypanosoma rangeli* in two localities of southern Ecuador identified *via* deep sequencing of mini-exon gene amplicons. Parasit Vectors. 2020;13: 252.

22. Bruschi F, Gradoni L (2018) The leishmaniases: old neglected tropical diseases. Cham, Switzerland: Springer. 245 pp. p.

23. Kato H, Gomez EA, Martini-Robles L, Muzzio J, Velez L, Calvopina M, et al. Geographic distribution of *Leishmania* species in Ecuador based on the *cytochrome B* gene sequence analysis. PLoS Negl Trop Dis. 2016;10: e0004844.

24. Bezemer JM, Freire-Paspuel BP, Schallig H, de Vries HJC, Calvopina M *Leishmania* species and clinical characteristics of Pacific and Amazon cutaneous leishmaniasis in Ecuador and determinants of health-seeking delay: a cross-sectional study. BMC Infect Dis. 2023;23: 395.

25. Hashiguchi Y, Velez LN, Villegas NV, Mimori T, Gomez EAL, Kato H Leishmaniases in Ecuador: comprehensive review and current status. Acta Trop. 2017;166: 299–315.

26. Arrivillaga-Henriquez J, Enriquez S, Romero V, Echeverria G, Perez-Barrera J, Poveda A, et al. [Eco-epidemiological aspects, natural detection and molecular identification of *Leishmania* spp. in *Lutzomyia reburra*, *Lutzomyia barrettoi majuscula* and *Lutzomyia trapidoi*]. Biomedica. 2017;37: 83–97.

27. Hashiguchi Y, Gomez EAL, Caceres AG, Velez LN, Villegas NV, Hashiguchi K, et al. Andean cutaneous leishmaniasis (Andean-CL, uta) in Peru and Ecuador: the causative Leishmania parasites and clinico-epidemiological features. Acta Trop. 2018;177: 135–145.

28. Magri A, Bianchi C, Chmelová L, Caffara M, Galuppi R, Fioravanti M, et al. Roe deer (*Capreolus capreolus*) are a novel potential reservoir for human visceral leishmaniasis in the Emilia-Romagna region of northeastern Italy. Int J Parasitol. 2022;52: 745–750.

29. Azami-Conesa I, Gomez-Munoz MT, Martinez-Diaz RA A systematic review (1990-2021) of wild animals infected with zoonotic *Leishmania*. Microorganisms. 2021;9: 1101.

30. Zhang JR, Guo XG, Chen H, Liu JL, Gong X, Chen DL, et al. Pathogenic *Leishmania* spp. detected in lizards from northwest China using molecular methods. BMC Vet Res. 2019;15: 446.

31. Rizzoli A, Tagliapietra V, Cagnacci F, Marini G, Arnoldi D, Rosso F, et al. Parasites and wildlife in a changing world: the vector-host- pathogen interaction as a learning case. Int J Parasitol Parasites Wildl. 2019;9: 394–401.

32. Kostygov A, Dobáková E, Grybchuk-Ieremenko A, Váhala D, Maslov DA, Votýpka J, et al. Novel trypanosomatid - bacterium association: evolution of endosymbiosis in action. mBio. 2016;7: e01985–01915.

33. Maslov DA, Westenberger SJ, Xu X, Campbell DA, Sturm NR Discovery and barcoding by analysis of spliced leader RNA gene sequences of new isolates of Trypanosomatidae from Heteroptera in Costa Rica and Ecuador. J Eukaryot Microbiol. 2007;54: 57–65.

34. Yurchenko V, Lukeš J, Tesařová M, Jirků M, Maslov DA Morphological discordance of the new trypanosomatid species phylogenetically associated with the genus *Crithidia*. Protist. 2008;159: 99–114.

35. Kozminsky E, Kraeva N, Ishemgulova A, Dobáková E, Lukeš J, Kment P, et al. Host-specificity of monoxenous trypanosomatids: statistical analysis of the distribution and transmission patterns of the parasites from Neotropical Heteroptera. Protist. 2015;166: 551–568.

36. Katakura K, Mimori T, Furuya M, Uezato H, Nonaka S, Okamoto M, et al. Identification of *Endotrypanum* species from a sloth, a squirrel and *Lutzomyia* sandflies in Ecuador by PCR amplification and sequencing of the mini-exon gene. J Vet Med Sci. 2003;65: 649–653.

37. Pinto CM, Ocana-Mayorga S, Tapia EE, Lobos SE, Zurita AP, Aguirre-Villacis F, et al. Bats, trypanosomes, and triatomines in Ecuador: new Insights into the diversity, transmission, and origins of *Trypanosoma cruzi* and Chagas disease. PLoS One. 2015;10: e0139999.

38. Medrano-Vizcaíno P, Grilo C, Brito-Zapata D, González-Suárez M Landscape and road features linked to wildlife mortality in the Amazon. Biodivers Conserv. 2023;32: 4337–4352.

39. Botero A, Thompson CK, Peacock CS, Clode PL, Nicholls PK, Wayne AF, et al. Trypanosomes genetic diversity, polyparasitism and the population decline of the critically endangered Australian marsupial, the brush tailed bettong or woylie (*Bettongia penicillata*). Int J Parasitol Parasites Wildl. 2013;2: 77–89.

40. McInnes LM, Gillett A, Ryan UM, Austen J, Campbell RS, Hanger J, et al. *Trypanosoma irwini* n. sp (Sarcomastigophora: Trypanosomatidae) from the koala (*Phascolarctos cinereus*). Parasitology. 2009;136: 875–885.

41. Maslov DA, Lukeš J, Jirků M, Simpson L Phylogeny of trypanosomes as inferred from the small and large subunit rRNAs: implications for the evolution of parasitism in the trypanosomatid protozoa. Mol Biochem Parasitol. 1996;75: 197–205.

42. Pereira A, Parreira R, Cristovao JM, Castelli G, Bruno F, Vitale F, et al. Phylogenetic insights on *Leishmania* detected in cats as revealed by nucleotide sequence analysis of multiple genetic markers. Infect Genet Evol. 2020;77: 104069.

43. García L, Ortiz S, Osorio G, Torrico MC, Torrico F, Solari A Phylogenetic analysis of Bolivian bat trypanosomes of the subgenus *Schizotrypanum* based on cytochrome B sequence and minicircle analyses. PLoS One. 2012;7: e36578.

44. Labruna MB, Whitworth T, Horta MC, Bouyer DH, McBride JW, Pinter A, et al. *Rickettsia* species infecting *Amblyomma cooperi* ticks from an area in the state of São Paulo, Brazil, where Brazilian spotted fever is endemic. J Clin Microbiol. 2004;42: 90–98.

45. Soares HS, Marcili A, Barbieri ARM, Minervino AHH, Moreira TR, Gennari SM, et al. Novel piroplasmid and *Hepatozoon* organisms infecting the wildlife of two regions of the Brazilian Amazon. Int J Parasitol Parasites Wildl. 2017;6: 115–121.

46. Shanmugasundram A, Starns D, Böhme U, Amos B, Wilkinson PA, Harb OS, et al. TriTrypDB: An integrated functional genomics resource for kinetoplastida. PLoS Negl Trop Dis. 2023;17: e0011058.

47. Klocek D, Grybchuk D, Macedo DH, Galan A, Votýpka J, Schmid-Hempel R, et al. RNA viruses of *Crithidia bombi*, a parasite of bumblebees. J Invertebr Pathol. 2023;201: 107991.

48. Katoh K, Standley DM MAFFT multiple sequence alignment software version 7: improvements in performance and usability. Mol Biol Evol. 2013;30: 772–780.

49. Wong TKF, Ly-Trong N, Ren H, Baños H, Roger AJ, Susko E, et al. IQ-TREE 3: phylogenomic inference software using complex evolutionary models. EcoEvoRxiv. 2025: 10.32942/x32942p32962n.

50. Ronquist F, Teslenko M, van der Mark P, Ayres DL, Darling A, Hohna S, et al. MrBayes 3.2: efficient Bayesian phylogenetic inference and model choice across a large model space. Syst Biol. 2012;61: 539–542.

51. Moreira D, López-García P, Vickerman K An updated view of kinetoplastid phylogeny using environmental sequences and a closer outgroup: proposal for a new classification of the class Kinetoplastea. Int J Syst Evol Microbiol. 2004;54: 1861–1875.

52. Albanaz ATS, Gerasimov ES, Shaw JJ, Sádlová J, Lukeš J, Volf P, et al. Genome analysis of *Endotrypanum* and *Porcisia* spp., closest phylogenetic relatives of *Leishmania*, highlights the role of amastins in shaping pathogenicity. Genes. 2021;12: 444.

53. Canizales-Silva C, Patiño LH, Jaimes-Dueñez J, Matta NE, Ramírez JD Characterization of novel *Trypanosoma* species from birds and amphibians in the Colombian dry forest using an integrative taxonomic approach. Int J Parasitol. 2026;(in press): 104830.

54. Nagy J, Fulmer AG, Löki V, Ruiz-Raya F, Hauber ME Biogeographic history, egg colouration, and habitat selection in *Turdus* thrushes (Aves: Turdidae). Biol Futur. 2023;74: 467–474.

55. Valkiūnas G, Iezhova TA, Carlson JS, Sehgal RN Two new *Trypanosoma* species from African birds, with notes on the taxonomy of avian trypanosomes. J Parasitol. 2011;97: 924–930.

56. Lima MB, Borges A, Wolf M, Santos HA, Dias RJP, Rossi MF First record of *Trypanosoma* (*Ornithotrypanum*) infecting Neotropical birds. Parasitol Res. 2024;123: 156.

57. Votýpka J, Zeman S, Stříbrná E, Pajer P, O. B, Kment P, et al. Multiple and frequent trypanosomatid co-infections of insects: the Cuban case study. Parasitology. 2024;151: 567–578.

58. Lukeš J, Tesařová M, Yurchenko V, Votýpka J Characterization of a new cosmopolitan genus of trypanosomatid parasites, Obscuromonas gen. nov. (Blastocrithidiinae subfam. nov.). Eur J Protistol. 2021;79: 125778.

59. Cleere N (1998) Nightjars: a guide to nightjars and related nightbirds. New Heaven, USA: Yale University Press. 317 p.

60. Cooke SC, Haskell LE, van Rees CB, Fessl B A review of the introduced smooth-billed ani *Crotophaga ani* in Galápagos. Biol Conserv. 2019;229: 38–49.

61. Lerner SDZ, Stauffer DF Habitat selection by blackburnian warblers wintering in Colombia. J Field Ornithol. 1998;69: 457–465.

62. Frolov AO, Kostygov AY, Yurchenko V Development of monoxenous trypanosomatids and phytomonads in insects. Trends Parasitol. 2021;37: 538–551.

63. Schulenberg TS, Stotz DF, Lane DF, O’Neill JP, Parker III TA (2010) Birds of Peru: revised and updated edition. Princeton, CT, USA: Princeton University Press. 664 p.

64. Camargo EP *Phytomonas* and other trypanosomatid parasites of plants and fruit. Adv Parasitol. 1999;42: 29–112.

65. von der Heyden S, Chao EE, Vickerman K, Cavalier-Smith T Ribosomal RNA phylogeny of bodonid and diplonemid flagellates and the evolution of euglenozoa. J Eukaryot Microbiol. 2004;51: 402–416.

66. Breunig A, König H, Brugerolle G, Vickerman K, Hertel H Isolation and ultrastructural features of a new strain of *Dimastigella trypaniformis* Sandon 1928 (Bodonina, Kinetoplastida) and comparison with a previously isolated strain. Eur J Protistol. 1993;29: 416–424.

67. Garcia MSA, Oliveira Filho VA, Brioschi MBC, Minori K, Miguel DC *Leishmania* (*Leishmania*) amazonensis. Trends Parasitol. 2025;41: 66–67.

68. Mendoza-Roldan JA, Latrofa MS, Tarallo VD, Manoj RR, Bezerra-Santos MA, Annoscia G, et al. *Leishmania* spp. in Squamata reptiles from the Mediterranean basin. Transbound Emerg Dis. 2022;69: 2856–2866.

69. Zhang JR, Guo XG, Liu JL, Zhou TH, Gong X, Chen DL, et al. Molecular detection, identification and phylogenetic inference of *Leishmania* spp. in some desert lizards from Northwest China by using internal transcribed spacer 1 (ITS1) sequences. Acta Trop. 2016;162: 83–94.

70. Otranto D, Testini G, Buonavoglia C, Parisi A, Brandonisio O, Circella E, et al. Experimental and field investigations on the role of birds as hosts of *Leishmania infantum*, with emphasis on the domestic chicken. Acta Trop. 2010;113: 80–83.

71. Matheus LMD, Duarte PO, de Ferreira EC First detection of *Leishmania* of the subgenus *Viannia* in *Alipiopsitta xanthops*, endemic bird of South America. Bioscience Journal. 2021;37.

72. Montaner-Angoiti E, Llobat L Is leishmaniasis the new emerging zoonosis in the world? Vet Res Commun. 2023;47: 1777–1799.

73. Alves MH, Mendoza-Roldan JA, Alfaro-Segura P, Carbonara M, Gomez A, Leitón NM, et al. Molecular detection of *Leishmania* and other vector-borne agents in free-ranging and captive herpetofauna from Costa Rica. Int J Parasitol Parasites Wildl. 2025;27: 101090.

74. Hong A, Zampieri RA, Shaw JJ, Floeter-Winter LM, Laranjeira-Silva MF One Health approach to leishmaniases: understanding the disease dynamics through diagnostic tools. Pathogens. 2020;9: 809.

75. Molyneux DH *Trypanosoma everetti* sp. nov. a trypanosome from the black-rumped waxbill *Estrilda t. troglodytes* Lichtenstein. Ann Trop Med Parasitol. 1973;67: 219–222.

76. Zídková L, Čepička I, Szabová J, Svobodová M Biodiversity of avian trypanosomes. Infect Genet Evol. 2012;12: 102–112.

77. Sereno D, Akhoundi M, Sayehmri K, Mirzaei A, Holzmuller P, Lejon V, et al. Noninvasive biological samples to detect and diagnose infections due to Trypanosomatidae parasites: a systematic review and meta-analysis. Int J Mol Sci. 2020;21: 1684.

78. Votýpka J, Pafčo B, Modrý D, Mbohli D, Tagg N, Petrželková KJ An unexpected diversity of trypanosomatids in fecal samples of great apes. Int J Parasitol Parasites Wildl. 2018;7: 322–325.

79. Zhang L, Liu S, Guo W, Lv C, Liu X Addressing biodiversity conservation, disease surveillance, and public health interventions through One Health approach in Hainan’s tropical rainforest. One Health Advances. 2024;2: 8.

